# Both the inflammatory response and clinical outcome differ markedly between adults with pneumococcal and meningococcal meningitis in a high HIV-1 prevalent setting in sub-Saharan Africa

**DOI:** 10.1101/539585

**Authors:** Emma C Wall, José Afonso Guerra-Assunção, Brigitte Denis, Matthew Scarborough, Katherine Ajdukiewicz, Katharine Cartwright, Mavuto Mukaka, Veronica S Mlozowa, Cristina Venturini, Theresa J Allain, David G Lalloo, Jeremy S Brown, Stephen B Gordon, Robert S Heyderman

## Abstract

Outcomes from pneumococcal meningitis (PM) are worse than meningococcal meningitis (MM), particularly in settings with high HIV-1 prevalence, but the reasons are unknown. We compared inflammatory responses between PM and MM in Malawian adults.

As compared to MM (n=27, 67% HIV-infected, mortality 11%), patients with PM (n=440, 84% HIV-infected, mortality 54%) were older, had strikingly lower CSF WCC, higher pro-inflammatory cytokine concentrations and higher mortality. PM is characterized by significantly lower CSF WCC, but greater inflammation and higher mortality compared to MM. Mechanistic understanding of blunting of the CSF leukocyte response in PM *in-vi*vo is required.

## Background

In most areas of the world, community-acquired acute bacterial meningitis (ABM) in adolescents and young adults is predominately caused by one of two bacterial pathogens, *Streptococcus pneumoniae* or *Neisseria meningitidis* [1, 2]. Although vaccination has impacted on ABM incidence in adults in well-resourced settings, in LMICs beyond the Sahel region in Africa, where both HIV infection and an intense force of bacterial carriage mean ABM remains a significant infectious disease [3, 4]. ABM caused by these pathogens is frequently indistinguishable at the bedside, but the clinical course is markedly different [5]. In LMICs case fatality rates (CFR) for pneumococcal meningitis (PM) reach 70%, compared to 5-10% for meningococcal meningitis (MM) [4, 6-9]. In all settings, 20-30% of survivors of PM experience complications including deafness, stroke and paralysis [2]. Estimates of years of life lost to disability (YLD) from PM are over twice those of MM [4, 10]. The two pathogens both have well-described host-pathogen interactions in animal models of meningitis and risk factors for acquisition in humans, but why outcomes differ markedly between PM and MM is not known [11-14].

Bacterial invasion into the CSF in ABM triggers a rapid-onset, localised inflammatory response, with resulting in migration of white cells across the blood-brain barrier to provide innate and adaptive immune responses in CSF to kill bacteria [15]. Both *S.pneumoniae* and *N.meningitidis* have differing mechanisms to avoid complement-mediated killing and phagocytosis in CSF [14, 16]. Of the small number of prospective clinical studies comparing the inflammatory response between the two pathogens, only complement activity [17, 18], and subtle differences between CSF cytokines [19-21] have been reported. To support development of new adjunctive treatments for ABM, we tested the hypothesis that differences in the cellular inflammatory response may differ in HIV-1 co-infected adults with ABM, accounting for differences in clinical outcomes in a high HIV prevalence setting in sub-Saharan Africa. We compared the inflammatory response in Malawian adults between cases of proven PM or MM using clinical, immunological and transcriptomic profiling.

## Methods

### Participants

Prospectively collected clinical data from adults and adolescents presenting to Queen Elizabeth Central Hospital in Blantyre, Malawi, with culture and PCR proven bacterial meningitis caused by either *S. pneumoniae* or *N. meningitidis* between 2003-2013 were extracted from the Malawi Meningitis Database (MMD) [22, 23]. CSF and blood samples were collected prior to administration of parenteral ceftriaxone. Clinical data were recorded on admission to hospital, repeat LP was performed for therapeutic monitoring at 48 hours in most participants recruited to clinical trials [24, 25]. Mortality was reported to 6 weeks [23].

### Procedures

Routine CSF microscopy, cell count, and CSF culture was done at the Malawi-Liverpool-Wellcome Trust Clinical Research Programme laboratory in Blantyre, Malawi as previously described [23]. Culture negative samples were screened using a multiplex PCR for *S. pneumoniae, N. meningitidis* and *Haemophilus influenzae type b* (Hib) (Fast-Track Diagnostics, Luxemburg). CSF cytokines were measured by bead array, as described [26]. CSF and whole blood for transcriptional profiling were collected in blood PAX-gene® (Pre-AnalytiX, Qiagen, USA) tubes, incubated for 4 hours at room temperature, and stored at −80 degrees°C. All patients were HIV tested using point-of care Genie™ HIV1&2 test kits (BioRad, USA).

RNA was extracted from blood and CSF using the PAXgene® Blood miRNA kit (Pre-Analytix, Qiagen, USA), quantified and RIN scores calculated using RNA Tapestation 4200® (Agilent, USA) and Nanodrop® (Thermoscientific, USA). Extracted RNA samples with RIN >7 underwent library preparation for polyA tailed mRNA with a RNA concentration of >1ng/1ul using with Kapa RNA hyperPrep kit (Roche), followed by 75 cycles of Next-generation sequencing with NextSeq® (Illumina®, USA).

### Bioinformatics and statistical analysis

Alpha <0.05 denoted statistical significance for all analyses, Mann-Whitney U tests were used on skewed data, and student’s *t* test for normally distributed data. 95% confidence intervals were presented for Odds Ratios. Logistic regression between clinical outcomes and risk factors was used to model associations while controlling for confounding factors. 95% confidence intervals presented for Odds Ratios. Fisher’s exact test was used to compare categorical variables. Graphical correlations were analysed using goodness of fit, statistical correlations calculated using Spearman’s test. Data were analysed in SPSS version 24 (IBM Statistics USA®) and GraphPad PRISM® version 7.0. Sequencing data quality was assessed using the FASTQC toolkit (http://www.bioinformatics.babraham.ac.uk/projects/fastqc/), sequenced cDNA libraries mapped at the transcript and gene levels to the human genome (assembly GRCh38) using *Salmon* v0.8.2 (http://salmon.readthedocs.io/en/latest/index.html).

The R package DESeq2 was used to test for differential gene expression. Data were expressed as log_2_ normalised gene counts, and FDR corrected p-values <0.05 used to test for significance. Network graphs of differentially expressed genes were generated using R package XGR (http://galahad.well.ox.ac.uk:3020/subneter/genes) and Gephi (https://gephi.org/),.

### Ethics

All participants or nominated guardians gave written informed consent for inclusion in the participating studies. Ethical approval for the transcriptomics study was granted by the College of Medicine Research and Ethics Committee (COMREC), University of Malawi, (P.01/10/980, January 2011) and the Liverpool School of Tropical Medicine Research Ethics Committee, UK (P10.70, November 2010), Liverpool, UK.

## Results

### Clinical data

We compared clinical details of proven PM cases (n=440) and MM cases (n=27) in the MMD (*Supplementary* Figure 1). Patients with PM were slightly older (OR 1.01 (95% CI 1.0 – 1.21) p<0.001), more likely to be female (OR 0.24 (0.09 : 0.6) p<0.01), living with HIV co-infection (OR 2.7 (1.1 – 6.5) p=0.02), anaemic (OR per 10mg/dL change 0.79 (0.69 : 0.90) p=0.001), have more severe clinical disease (OR 1.57, 95% CI 1.01-2.4 p=0.04), and a substantially higher case fatality rate (CFR) (OR 8.5 (2.5-28.1) p<0.001) (Table 1). After multivariate correction, the strongest variables that discriminated PM from MM were mortality rate, age, gender and CSF WCC.

**Table 1:**
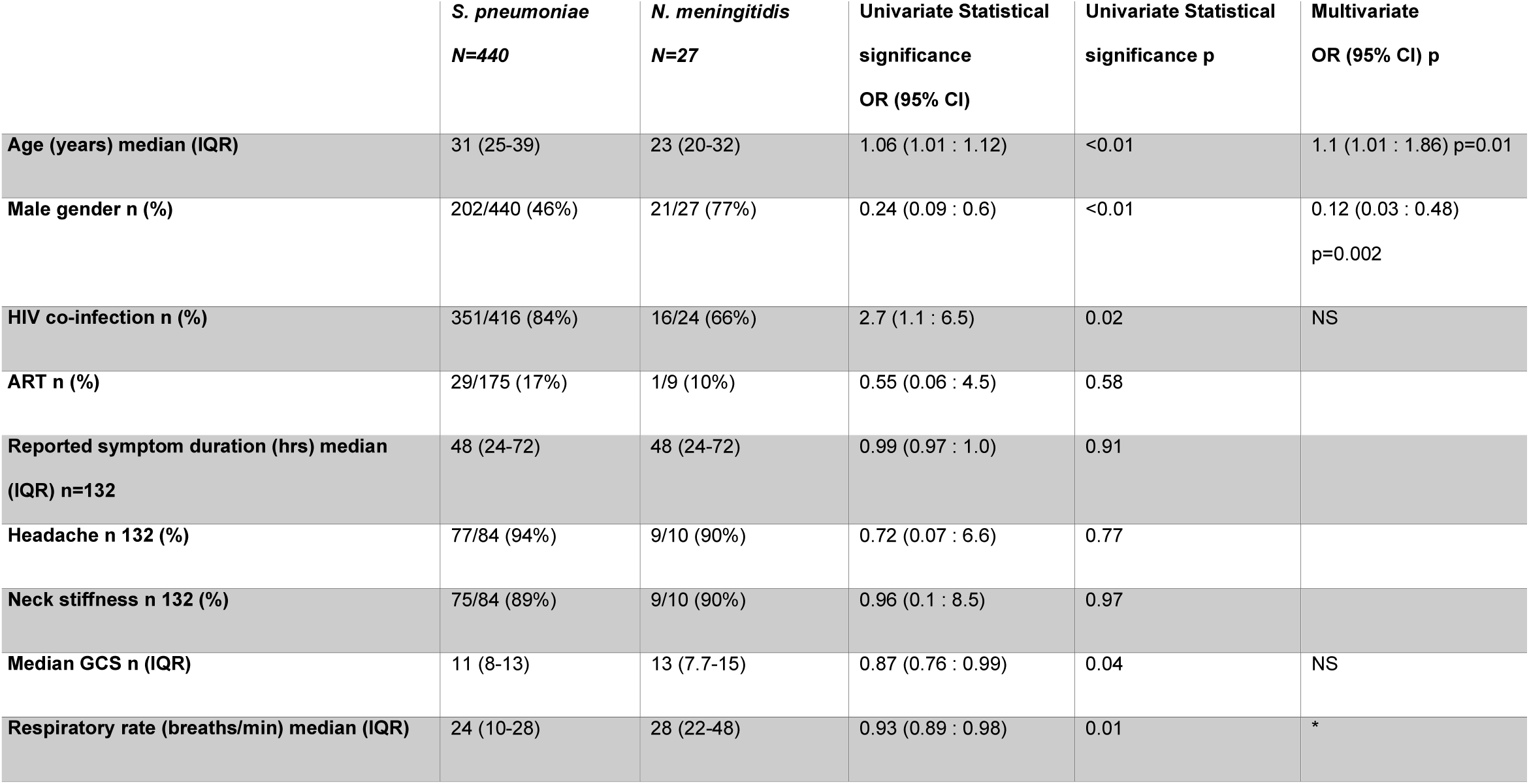
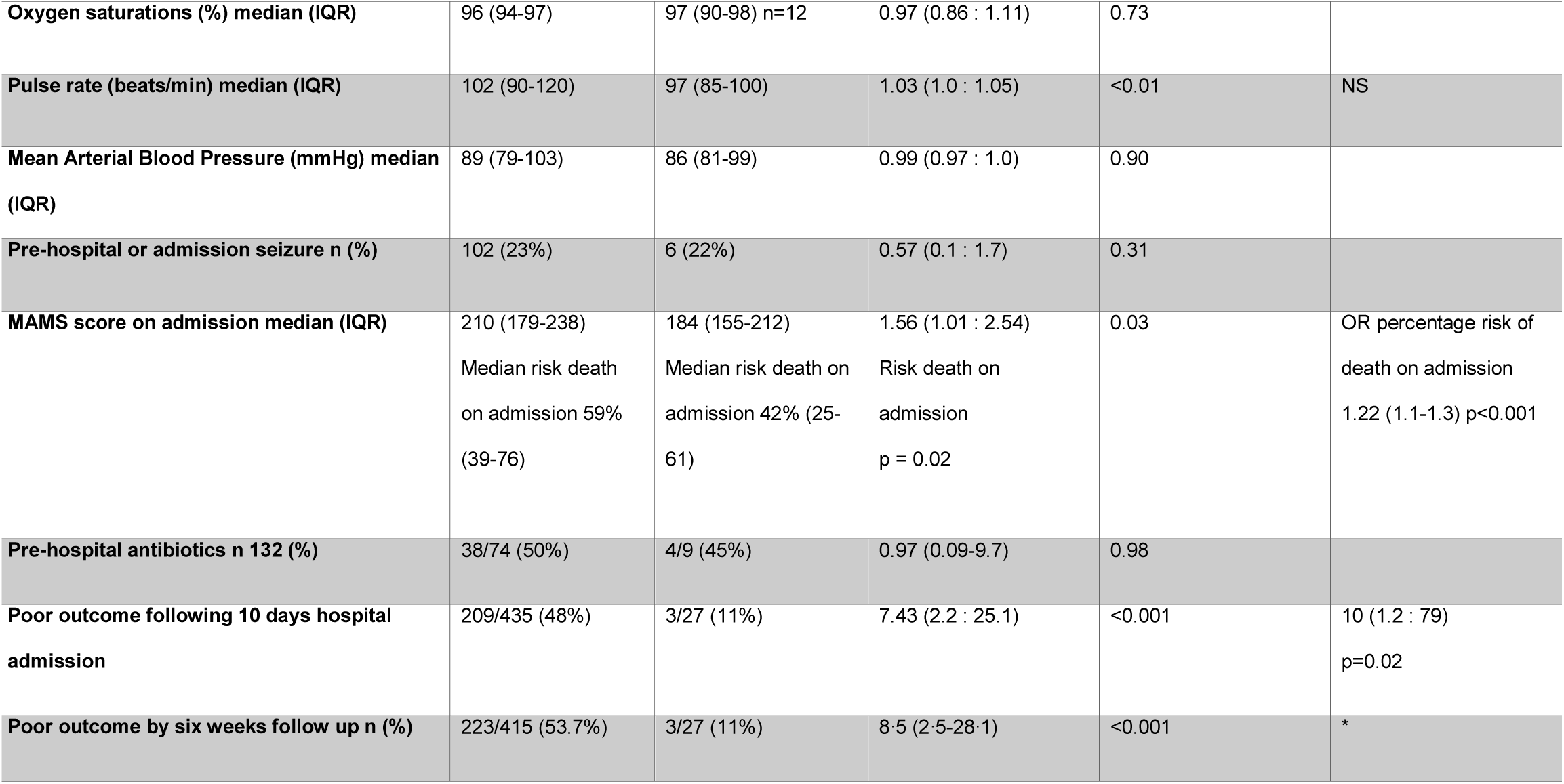
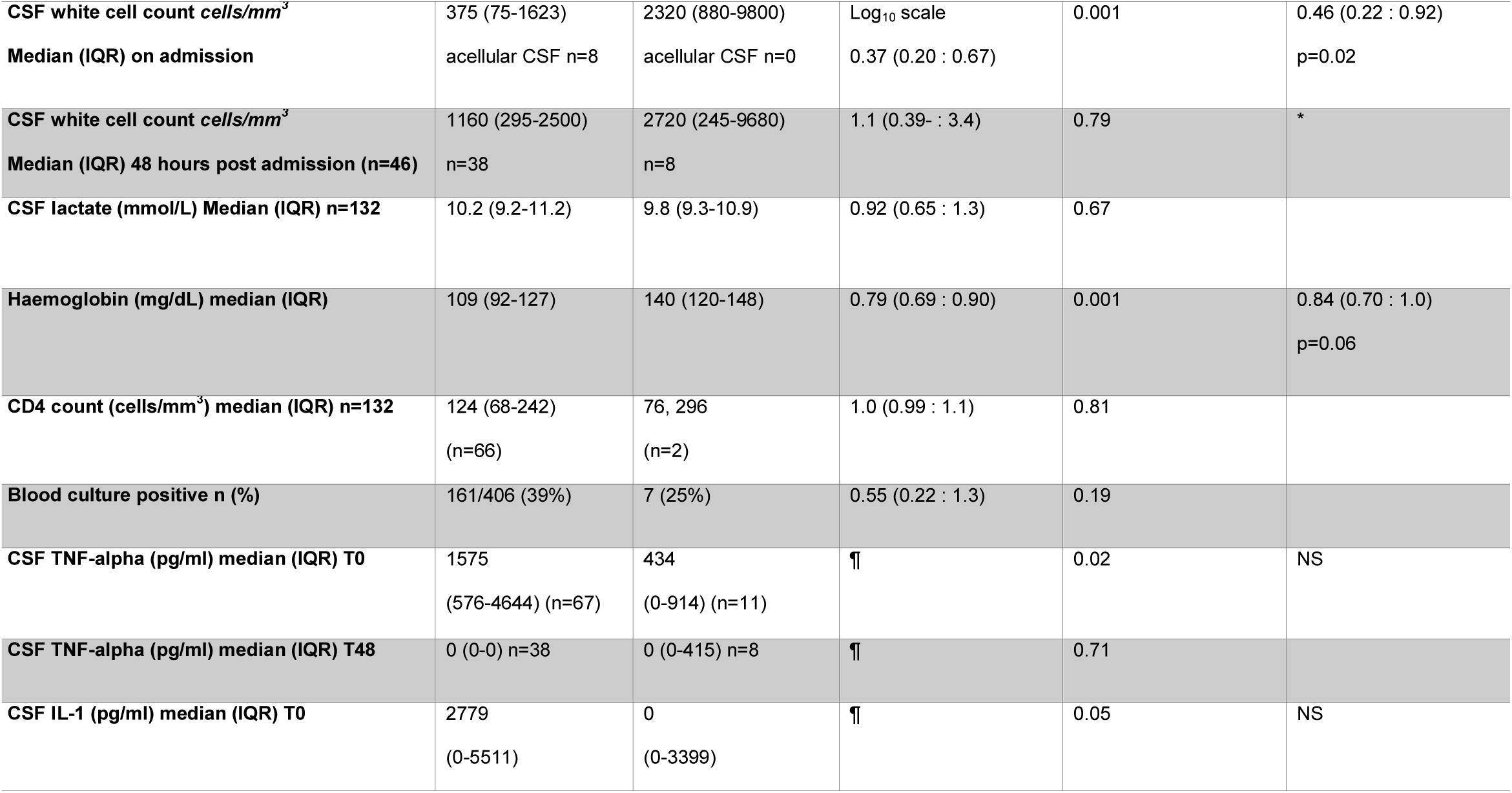
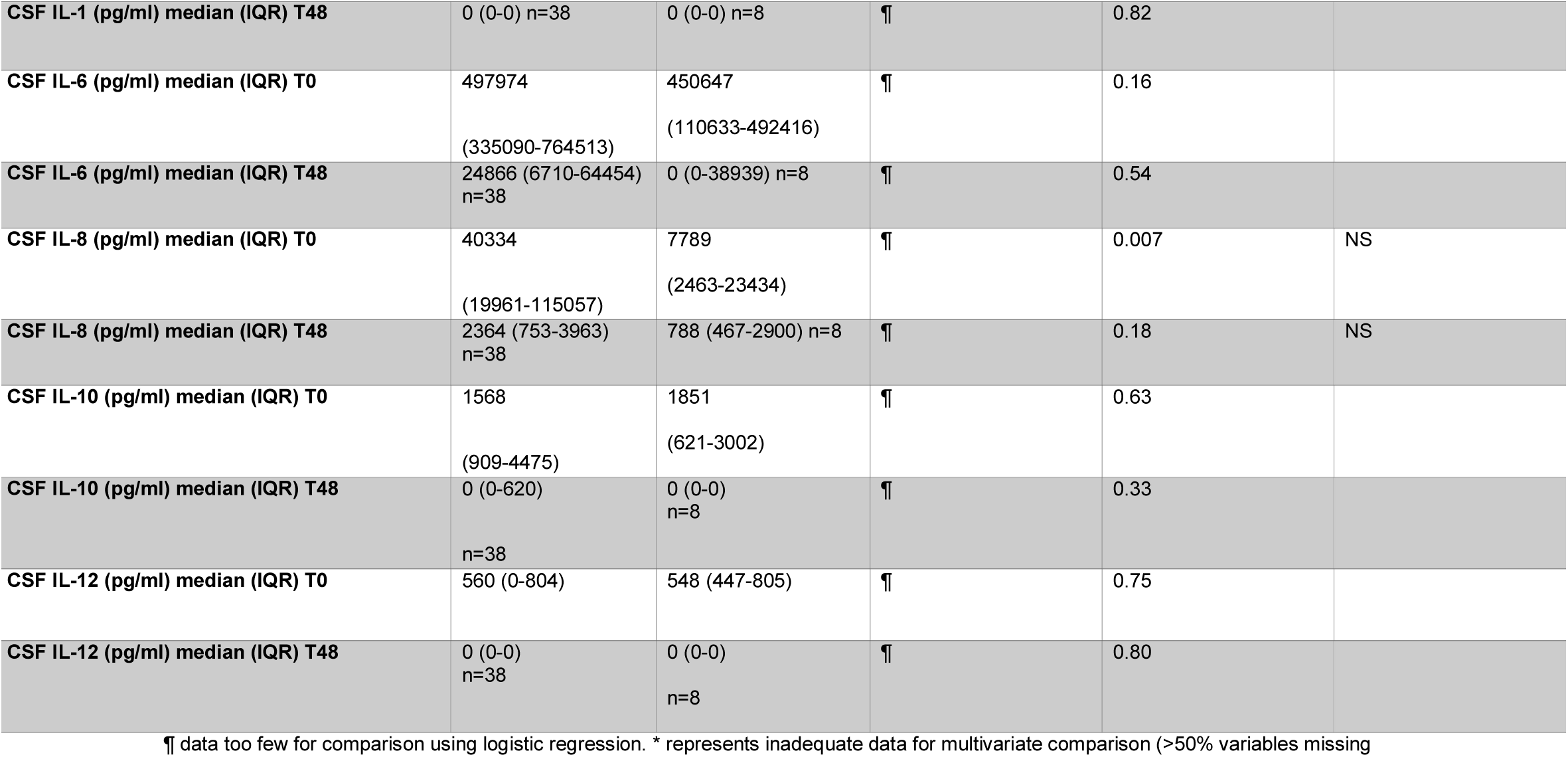
Demographic details of included patients comparing cases of proven bacterial meningitis caused by *Streptococcus pneumoniae* and *Neisseria meningitidis*. Clinical and laboratory characteristics of adults and adolescents with pneumococcal and meningococcal meningitis

### CSF inflammatory profiling

Median CSF WCC (>50% neutrophils) were lower in cases of PM than MM (adjusted OR 0.46 (0.22 : 0.92, p=0.02) (Figure 1A) (Table 1). Median CSF WCC in PM were low on admission; 375 cells/mm^3^ (IQR 75-1623), incrementing over 48 hours to 1160 cells/mm^3^ (IQR 295-2500) p<0.001. CSF WCC in MM were significantly higher on admission (2320 cells/mm^3^ (IQR 880-9800) and at 48 hours; 2720 cells/mm^3^ (IQR 245-9680) p=0.34 (Figure 1A).

**Figure 1:**
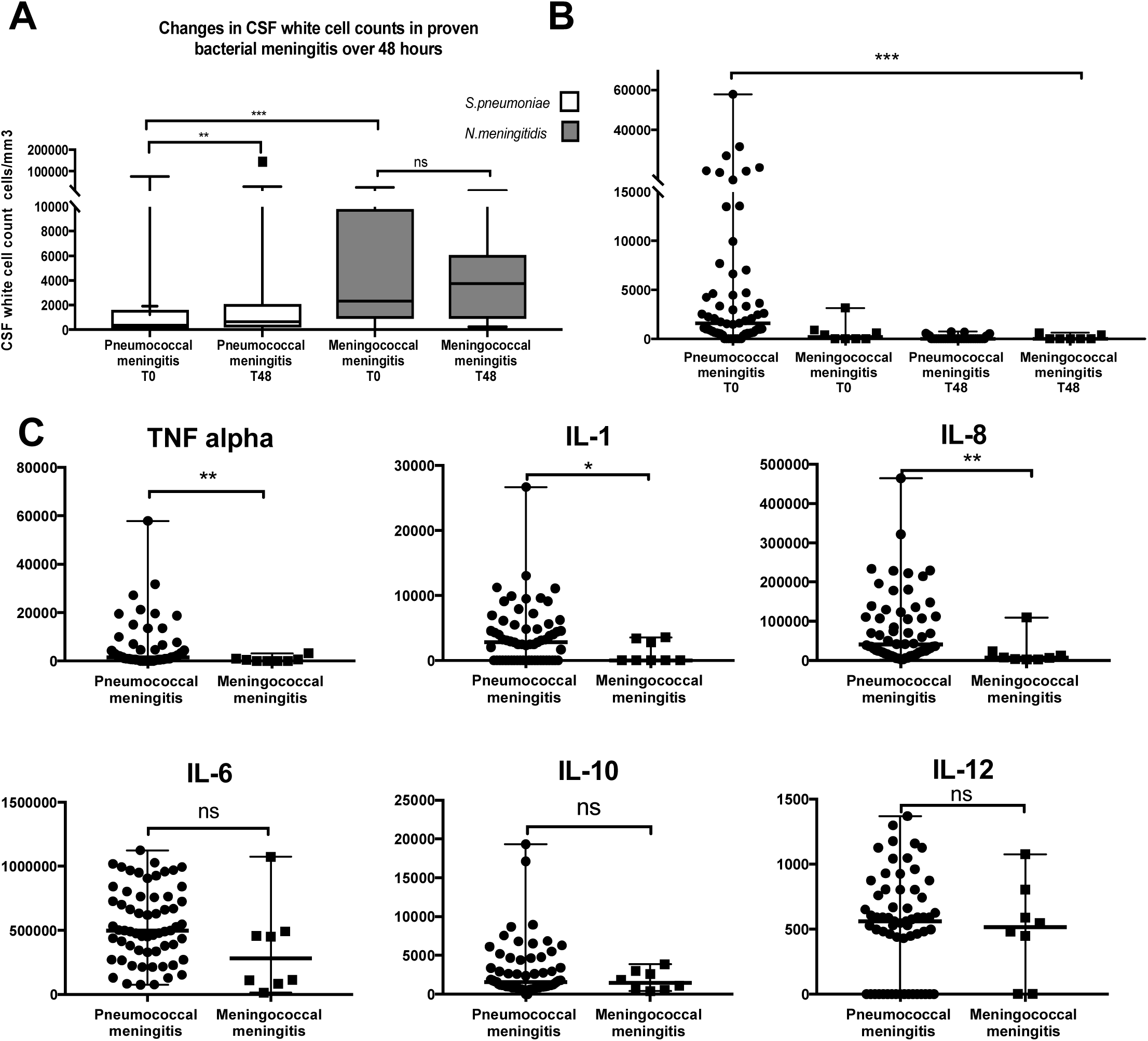
Inflammatory profiling of cases of pneumococcal meningitis and meningococcal meningitis reveals patients with PM have significantly lower CSF white cell counts and significantly higher CSF pro-inflammatory cytokines. CSF white cell count (WCC) on admission to hospital is statistically significantly lower in pneumococcal meningitis (PM) compared to meningococcal meningitis (MM), rising in PM after 48 hours but not in MM (A). Comparison of CSF white cell count (cells/mm^3^) between cases of proven pneumococcal meningitis (PM, n=440, white) and meningococcal meningitis (MM, n=27, shaded grey) on admission. Y axis CSF WCC in cells/mm^3^, boxes indicate interquartile range, bars indicate median with range. CSF TNF alpha production dramatically falls in pneumococcal meningitis after 48 hours of antibiotic therapy (B). Comparison of CSF cytokine levels on admission between pneumococcal and meningococcal meningitis measured by bead array, represented on Y axis in pg/uL, individual samples represented by single dot (PM, n=67) or square (MM, n=8) at on admission T0, or after 48 hours of treatment (T48) Key pro-inflammatory cytokines are higher in the CSF in cases of PM than MM on admission to hospital (C). CSF cytokines at time 0 (Y axis). All bars indicate median with range. Analyses done using Mann-Whitney U test (A-C) and Kruskall-Wallis Test (B). p<0.05 determined significance.*=p<0.05, **=p<0.01, *** p<0.001

In a sub-set of patients who had CSF cytokine profiling, TNF alpha, IL-8 (CXCL8) and IL-1 in CSF were significantly higher from cases of PM (n=66) compared to MM (n=11) on admission to hospital (Figure 1C) (Table 1). All CSF cytokines were lower at 48 hours (Figure 1B). CSF WCC did not correlate with any of tested CSF cytokines, including IL-8 (CXCL8) (*S.*Figure 2A&B). There was little evidence of systemic cytokine responses in blood (*S*.Figure 3). Ratios of median CSF: blood cytokine levels (pg/ul) were: TNF-a (124:1), IL-8 (321:1), IL-6 (365:1), and IL-10 (62:1). Data on MM were too few to calculate CSF: Blood cytokine ratios.

### The role of HIV-1 infection

Case Fatality Rates (CFR) from PM and MM did not differ by HIV-1 infection status; of patients with PM, 352/417 (84%) were HIV co-infected, CFR was 179/335 (53%) in HIV co-infected patients compared to 33/61 (54%) in HIV negative, OR 0.97 (95% CI 0.57 – 1.6) p=0.91. Of patients with MM whose HIV-1 status was known, 16/24 (67%) were HIV co-infected, CFR was 0/16 in HIV co-infected and 1/8 (13%) in HIV negative p=0.33. HIV-1 co-infection had no appreciable effect on the CSF WCC response in either PM or MM (PM OR 1.0 CI 1.0-1.1, p=0.22); (MM OR 1.0 CI 1.0-1.01, p= 0.78). Median CD4 counts in HIV co-infected individuals were low (124 cells/mm^3^ IQR 68-242) (Table 1).

### CSF and blood transcriptomics

To examine the wider inflammatory response using transcriptomics between cases of PM and MM, we undertook paired CSF and blood transcriptional profiling using RNAseq (Supplementary Figure 1) from paired blood and CSF samples of thirty-two adults with proven ABM prior to antibiotic administration (PM n=28, MM n=4). Using DESeq2, we tested for differential gene expression in CSF and blood. Although the number of MM samples were limited, the results show that differentially expressed genes between PM and MM were more marked in CSF than in blood (*S*. Figure 4A&B). Differentially expressed genes were tested for weighted correlation networks using XGR, co-correlated genes were then analysed by network centrality analysis using Gephi. Upregulated genes in in MM centered around genes coding for neutrophil recruitment (*CXCL8, CD44, PIK3CG*), endothelial activation (*VAS3, PTGS2*) and cellular signalling and proliferation (*KRAS, CASR*) (*S*.Figure 4C). In contrast, upregulated genes in PM centred around cellular transcriptional (*HRAS, ABCC1, RAD50*) metabolic (*SUCLA2, FASN, NME3*) and non-specific inflammatory processes (*S*.Figure 4D).

## Discussion

We report strikingly different acute inflammatory responses in CSF between meningitis due to the two common community-acquired pathogens *S.pneumoniae* and *N.meningitidis* in predominately HIV-1 infected adults in Malawi. Patients with PM, paradoxically, had significant blunting of CSF white cell counts CSF compared to patients with MM, but a contrastingly more intense pro-inflammatory cytokine response, irrespective of HIV-1 serostatus. Although the host-pathogen interaction in the immunologically protected CSF space in ABM is well-recognised to be highly pro-inflammatory [15], our study documents differing inflammatory responses that are likely to require different anti-inflammatory approaches to improve clinical outcomes. Thus, our study has important implications for the development of better adjuncts to antibiotics in our setting.

Patients with PM in our study had CSF WCC that were 84% lower and were 8 times more likely to die than patients with MM. Whilst low CSF WCC are universally associated with poor outcomes from ABM, [7, 12] our study is the first to describe such a significant difference in CSF cell counts between these two common causes of ABM in a LMIC setting. The underlying mechanisms causing CSF leukopaenia in ABM are poorly understood. In animal models, neutrophil destruction by *S.pneumoniae* cytolysins, including the TLR-4 agonist pneumolysin has been proposed as a cause of CSF leukopaenia in PM [27, 28]. We have previously reported an association between persistence of pneumolysin and outcome in our setting, independent of bacterial load, suggesting pneumolysin may be an important component of the host-pathogen interaction in PM, but mechanistic data in humans are lacking [26, 29]. In contrast, *N.meningitidis* is not known to synthesise similarly destructive cytolysins, instead surviving primarily by evading complement-mediated killing [30]. Patients with MM in our setting clinically present with significantly greater CSF innate responses, with substantially better outcomes. Understanding the causes of CSF WCC blunting *in-vivo*, particularly the role of cytolysins, will be essential to improve outcomes, particularly from PM.

Although CSF pro-inflammatory cytokines were higher in PM, however the absence of reported correlation between CSF WCC and cytokines in either PM or MM was surprising. In animal models of PM, blocking neutrophil migration to the CSF compartment reduces CSF cytokine production, suggesting neutrophils produce pro-inflammatory cytokines in CSF [31, 32]. In our patients, we hypothese that these proteins may either be released from necrotic, and thus uncounted leukocytes in the CSF, or from innate neuronal cells such as microglia as opposed to neutrophils alone [27, 33, 34]. Further studies to investigate the cellular source of CSF cytokines *in-vivo* in both PM and MM are required.

Most of our patients were HIV-1 co-infected, interestingly neither outcome nor CSF WCC differed between those with or without HIV co-infection. HIV is a well-recognised risk factor for acquisition of both PM and MM [35, 36]. However, HIV appears to have little impact on the established inflammatory response in either infection, although the proportion of HIV-negative patients was relatively small. HIV-1 is not associated with clinical outcome from PM, although it may be a risk factor for mortality in MM [7, 37, 38]. We conclude that high CFR from ABM in our setting is not strongly associated with HIV-1.

Our study was limited by lack of stratification of the inflammatory profiles by bacterial load, CD4 count and HIV-1 viral load; these data were not routinely collected during the clinical trials. Similarly, the CSF cytokine data were limited by the volume of CSF available in some cases. CSF transcriptome data were only available for 2 patients with MM, so although we reported significant differential gene expression in CSF, these results must be considered preliminary. Furthermore, we could not stratify the transcriptome results by neutrophil count. Finally, the data were obtained from patients in a high disease burden, low income setting with high HIV prevalence, and may not be generalisable to other settings.

In summary, our study demonstrates that despite higher CSF pro-inflammatory cytokine levels, patients with PM have a paradoxically blunted CSF WCC and worse clinical outcomes compared to cases of MM. To improve outcomes from ABM, further dissection of the *in-vivo* host-pathogen interactions in PM and MM, to understand the mechanisms causing blunting of the acute CSF WCC response in PM is required.

## Funding

This study was funded by a Clinical Lecturer Starter Grant from the Academy of Medical Sciences (UK), a PhD Fellowship in Global Health to EW (089671/B/09/Z) and a Career Development Fellowship to SG (2005) from the Wellcome Trust. CSF molecular diagnostics were funded by a Royal Society of Tropical Medicine and Hygiene small grants for Africa award to EW. The SAM trial was supported by the Meningitis Research Foundation, and the Malawi-Liverpool-Wellcome Trust Clinical Research Programme is supported by core funding from the Wellcome Trust. This work was undertaken at UCLH/UCL who received a proportion of funding from the Department of Health’s NIHR Biomedical Research Centre’s funding scheme. The funders of the study had no role in study design, data collection, data analysis, data interpretation, or writing of the report. The corresponding author had full access to all the data and the final responsibility to submit for publication.

## Acknowledgements

The authors would like to thank the study patients and guardians, the Bundles for Adult Meningitis (BAM) research team, clinical and laboratory staff at the Queen Elizabeth Central Hospital and Malawi-Liverpool-Wellcome Trust Clinical Research Programme in Blantyre Malawi for support given during the study and Professor Mike Levin and Dr Victoria Wright of Imperial College UK for support at the early stages of the project and gift of PAXgene tubes for RNA storage. We would like to thank Professor Judy Breuer and the staff of the Pathogen Genomics Unit at University College London for their assistance with library preparation and RNA sequencing.

## Supplementary data

### RNAseq repository of all CSF and blood data

Mapped, sequence files for all included patients are available on a consent-basis through the European Phenome-Genome Archive at the European Bioinformatics Institute (EBI) https://www.ebi.ac.uk/ega/studies/EGAS00001003355

**S. Figure 1:**
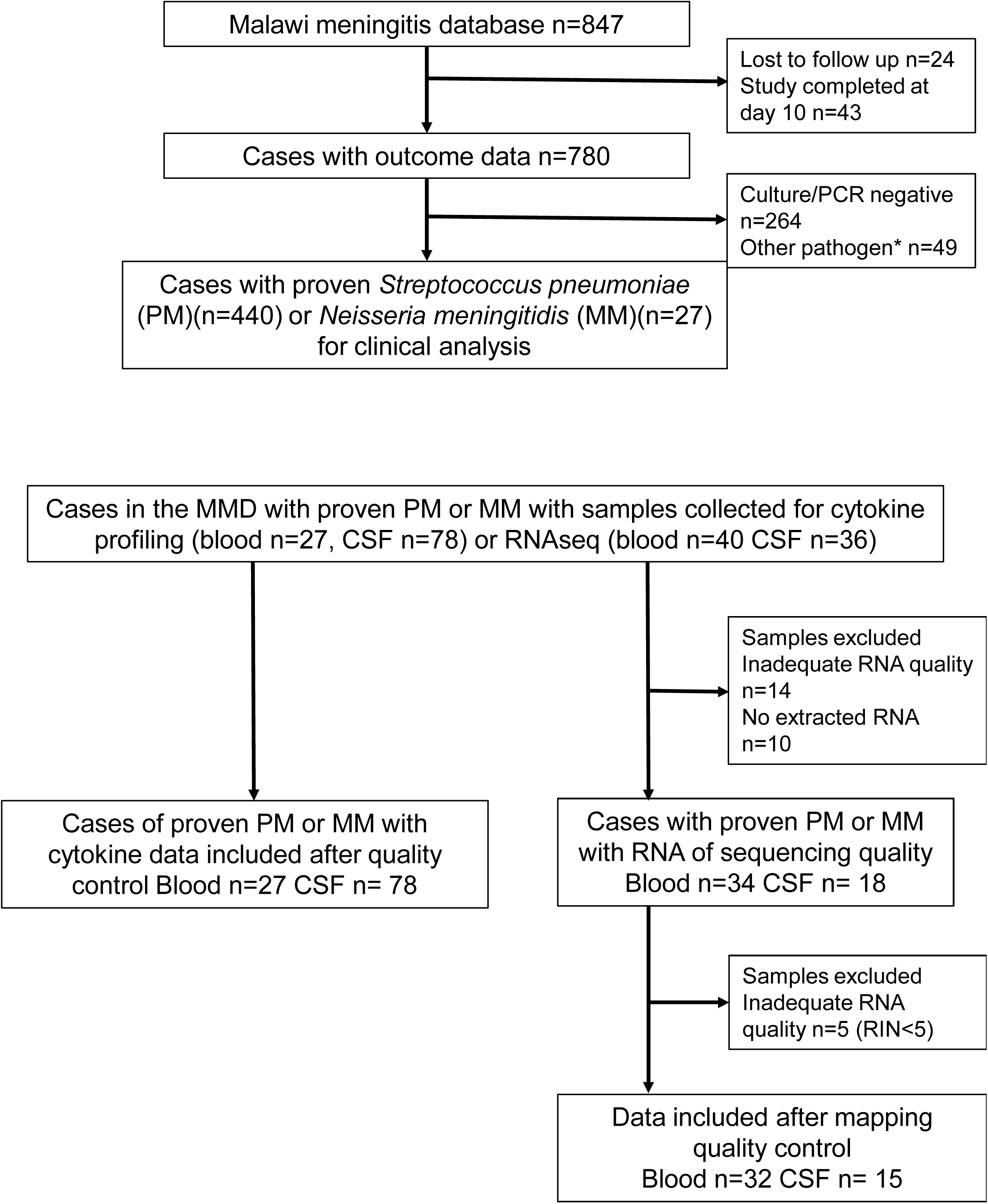
Selection of study patients for inclusion

**S. Figure 2:**
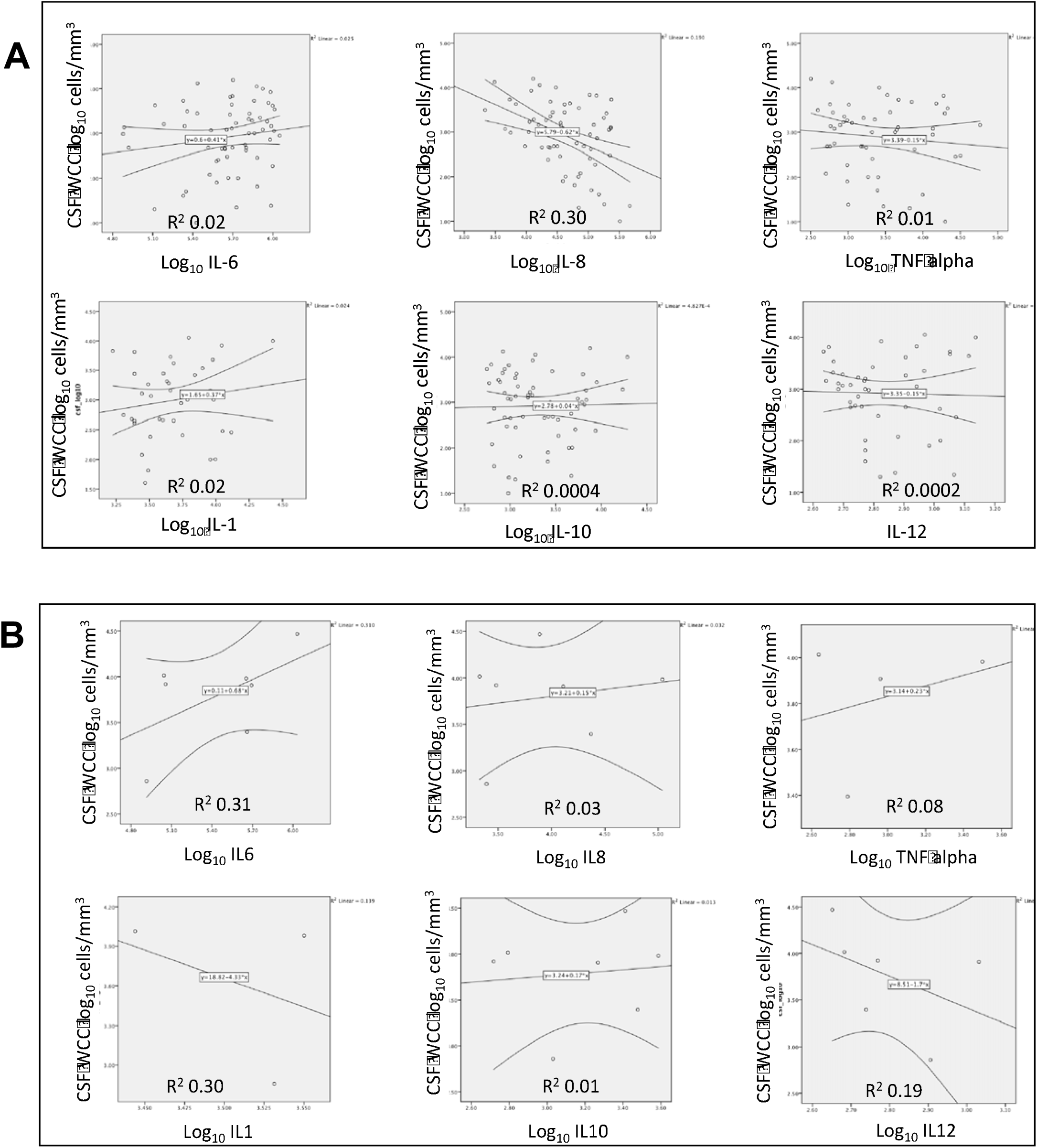
CSF white cell counts and individual CSF cytokine levels do not correlate in either cases of pneumococcal meningitis (A) or meningococcal meningitis (B). Log_10_ levels of cytokine in the CSF (pg/ul) are shown on the X axis, log_10_ CSF WCC on the Y axis. Lines of best fit are shown with 95% confidence intervals (curved lines) calculated using Spearman’s correlation, R^2^ values are shown per plot.

**S. Figure 3:**
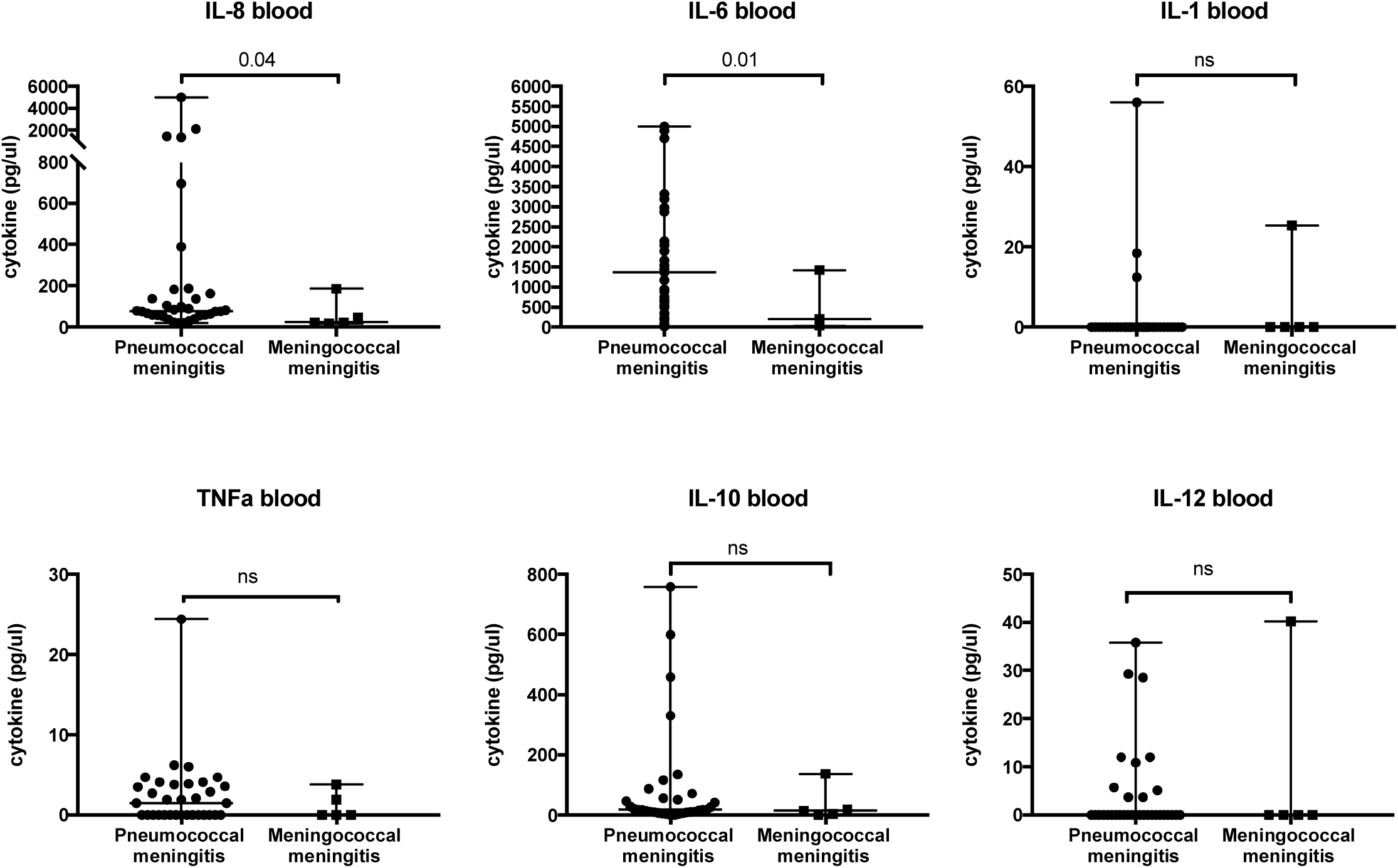
Cases of pneumococcal meningitis have correspondingly higher levels of IL-8 and IL-6 in the blood than cases of meningococcal meningitis. Comparison of cytokine levels in blood, on admission between pneumococcal and meningococcal meningitis, measured by bead array, represented on Y axis in pg/uL. Individual samples represented by single dot (PM, n=21) or square (MM, n=6) at on admission (T0), or after 48 hours of treatment (T48). All bars indicate median with range. Analyses done using Mann-Whitney U test. p<0.05 determined significance.

**S. Figure 4:**
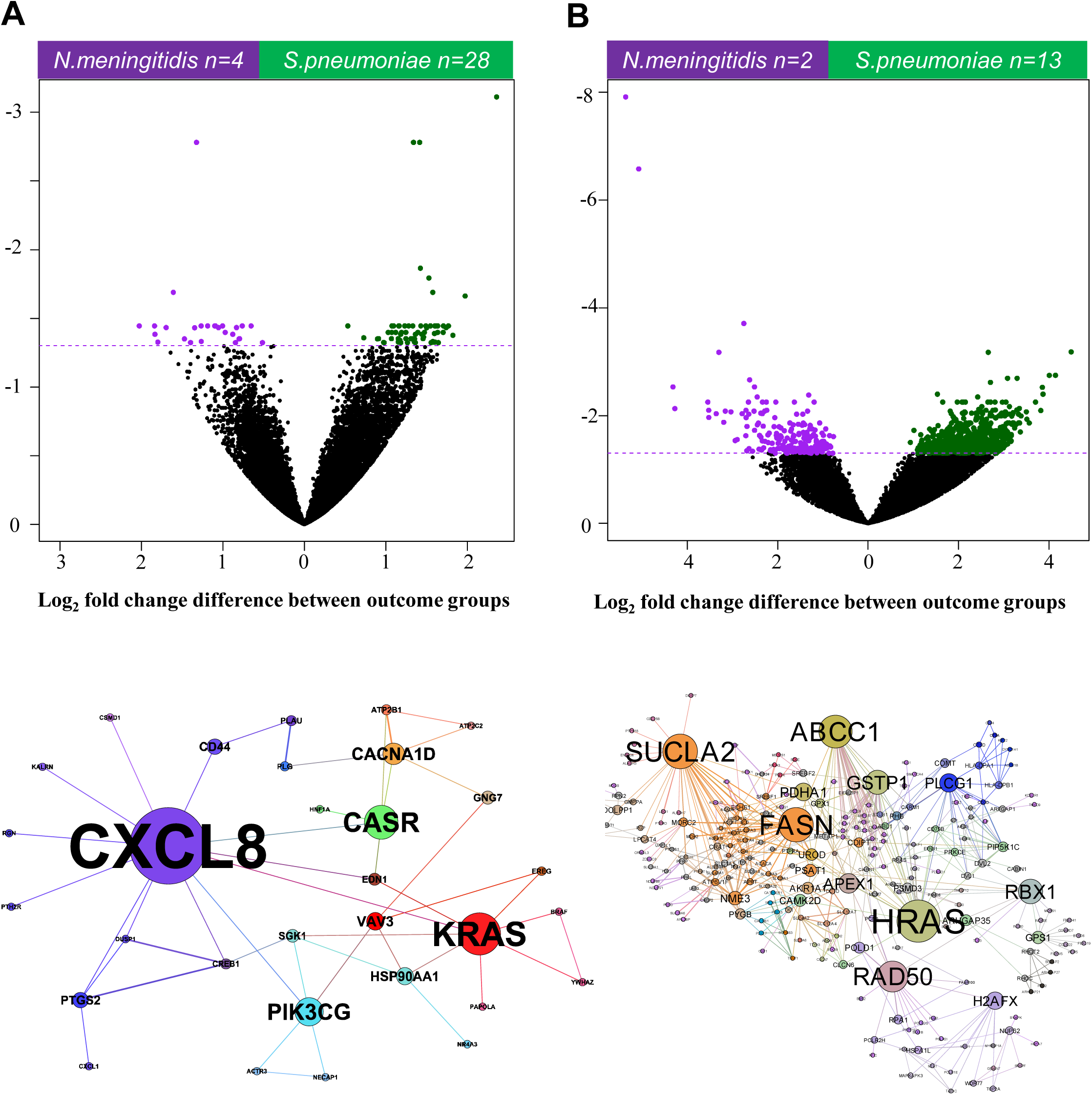
Transcriptional profiling with RNAseq reveals differential gene expression between pneumococcal and meningococcal meningitis in the CSF compartment. Volcano plot describing differential gene expression in Blood compartment (2A, n=32) and CSF (B, n=15) between *N.meningitidis* (upper left quadrant, purple) and *S.pneumoniae* (upper right quadrant, green) meningitis. Coloured dots represent individual genes expressed over 1 log_2_ fold change differential expression (X axis) with adjusted log_10_ False Discovery Rate (FDR) p value (Y axis) <0.05 (horizontal dashed line). Networks of differentially expressed genes in the CSF show upregulation of genes for neutrophil recruitment and endothelial activity in MM (2C) compared to house-keeping and metabolic gene expression in PM (2D). Labelled circles represent individual genes expressed over 1 log_2_ fold change differential expression with adjusted log_10_ False Discovery Rate (FDR) p value <0.05. Gene names are proportionally sized for network centrality and coloured for clusters as follows: Dark blue = innate and adaptive immunological network signaling, red = endothelial activation, light blue = signalling cascade activation, orange = metabolic activity, pink = DNA transcription, translation and repair, pale green = metabolic signaling, grey = ubiquitination of proteins. Networks created from differentially expressed genes in CSF using XGR, analysed by network centrality using Gephi.

